# βII-spectrin is required for synaptic positioning during retinal development

**DOI:** 10.1101/2022.12.07.519458

**Authors:** Debalina Goswami-Sewell, Caitlin Bagnetto, Joseph T Anderson, Akash Maheshwari, Elizabeth Zuniga-Sanchez

## Abstract

Neural circuit assembly is a multi-step process where synaptic partners are often born at distinct developmental stages, and yet they must find each other and form precise synaptic connections with one another. This developmental process often relies on late-born neurons extending their processes to the appropriate layer to find and make synaptic connections to their early-born targets. The molecular mechanism responsible for the integration of late-born neurons into an emerging neural circuit remains unclear. Here we uncovered a new role for the cytoskeletal protein βII-spectrin in properly positioning pre- and post-synaptic neurons to the developing synaptic layer. Loss of βII-spectrin disrupts retinal lamination, leads to synaptic connectivity defects, and results in impaired visual function. Together, these findings highlight a new function of βII-spectrin in assembling neural circuits in the mouse outer retina.

**Highlights:**

- Established a new role for βII-spectrin in assembling retinal circuits
- βII-spectrin positions pre- and post-synaptic neurons to the developing synaptic layer
- Early positioning of processes to the OPL is required for synaptogenesis
- Loss of βII-spectrin disrupts synaptic connectivity and impairs visual function

## INTRODUCTION

During the development of the central nervous system, different neuron subtypes that assemble into a functional circuit are often born at various developmental time points (Faux et al., 2012). In order to find their respective synaptic partner, late-born neurons have the daunting task of extending their dendrites and axon into the correct layer (Faux et al., 2012). However, the molecular mechanism of how late-born neurons know where to precisely extend their processes to form synapses with their respective targets remains poorly understood.

The mammalian retina, which is an extension of the central nervous system, is an excellent model to study this question. Within the retina, neurons are born at different developmental time points (Young, 1985), and yet they find their respective synaptic partner with a high degree of specificity. In the outer retina, cone photoreceptors synapse selectively to the dendrites of cone bipolars and the dendrites of horizontal cells, whereas rod photoreceptors synapse with the dendrites of rod bipolars and the axon terminal of horizontal cells (**Figure 1A**). Synaptic connections are confined to the outer plexiform layer (OPL) and inner plexiform layer (IPL), whereas cell bodies of photoreceptors, interneurons, and ganglion cells are localized to the outer nuclear layer (ONL), inner nuclear layer (INL), and ganglion cell layer (GCL). See **Figure 1A**. During development, cone photoreceptors first extend their axon terminal and make contacts to horizontal cells in the presumptive OPL, which is marked by the separation between the ONL and INL (Burger et al., 2021; Sarin et al., 2018) as shown in **Figure 1B**. Early OPL formation is observed starting at postnatal day (P) 5 (Burger et al., 2021; Sarin et al., 2018). Bipolar neurons are born between P0-3 (Carter-Dawson & LaVail, 1979; Young, 1985) but do not extend their dendrites into the OPL until P5 (Morgan et al., 2006). Live imaging of bipolar neurons reveal that dendrites emerge from a long neuroepithelial-like process that spans the entire retina (Morgan et al., 2006). Interestingly, only the dendrites of bipolars found within the developing OPL are stabilized and maintained whereas those that emerge in other regions are rapidly destabilized and eliminated (Morgan et al., 2006). See **Figure 1B**. From P9-13, the long neuroepithelial-like processes from bipolar neurons disappear (Morgan et al., 2006), and synapses between photoreceptors and bipolar neurons form (Sarin et al., 2018). We refer to this timepoint as a midpoint of OPL formation as depicted in **Figure 1B**. By P21, synapse formation in the outer retina is largely complete (Sarin et al., 2018), which we refer to as the end of OPL formation in **Figure 1B**.

**Figure 1:**
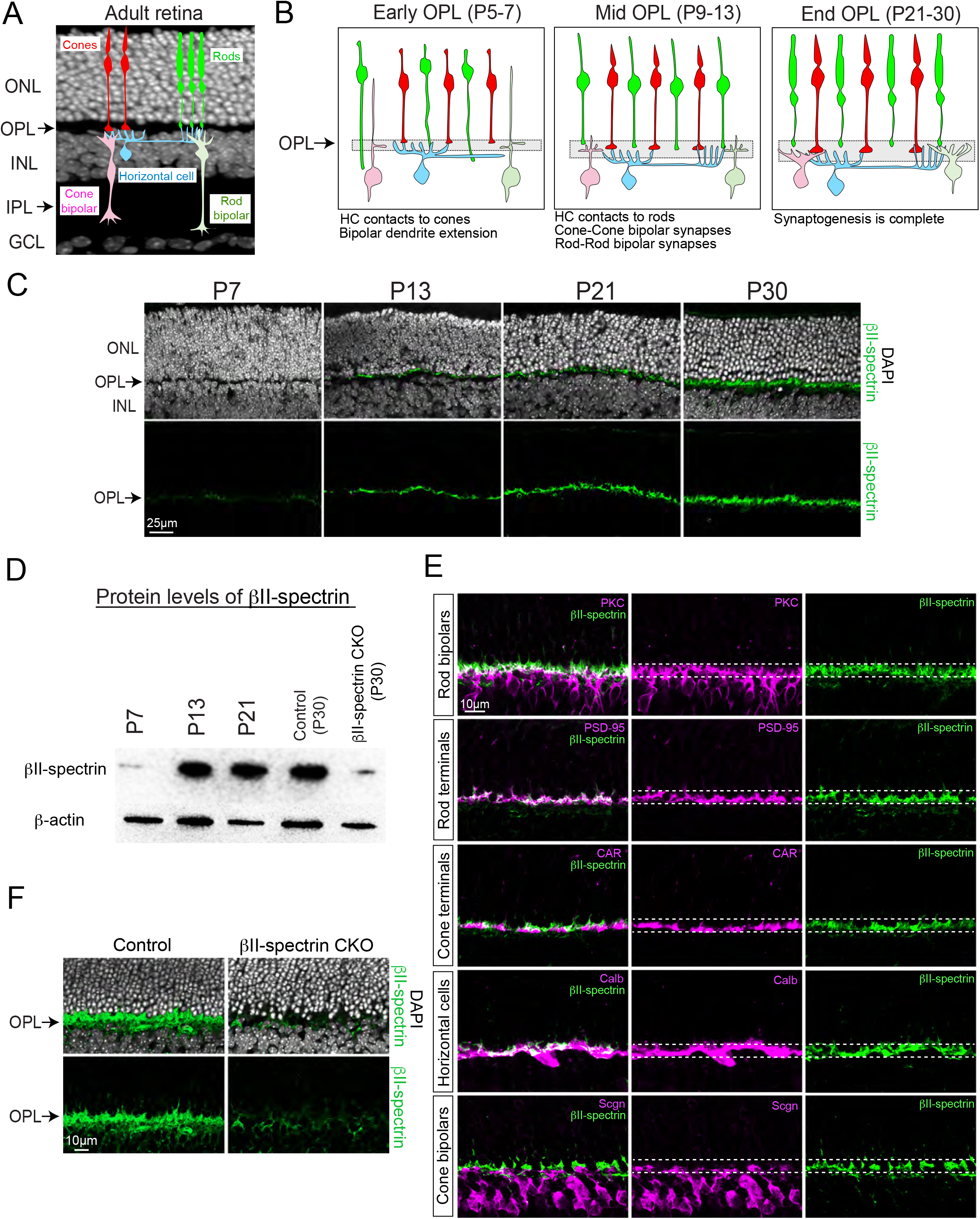
βII-spectrin is localized to the OPL during synaptogenesis (A-F). Schematic drawing of the adult retina. The retina is subdivided into different three nuclear layers (i.e. ONL, INL, GCL) and two synaptic layers (i.e. OPL, IPL) (A). Cone photoreceptors (red) synapse to the dendrites of horizontal cells (light blue) and dendrites of cone bipolars (light red). Rod photoreceptors (green) synapse to the axon terminal of horizontal cells (light blue) and the dendrites of rod bipolars (light green) (A). Development of the OPL (B). Horizontal cells (light blue) first make contacts to cones (red) in the presumptive OPL where cone bipolars (light red) and rod bipolars (light green) begin to extend their dendrites. From P9-13, bipolar neurons continue to extend their dendrites to the OPL and make synaptic connections to their targets. By P21, synapse formation in the retina is largely complete (B). Antibody staining of βII-spectrin (green) in wild-type retinas shows protein localization in the OPL that gradually increases from P7 to P30 (C). Nuclei is stained with DAPI (C). Western blot analysis reveals βII-spectrin protein levels increase from P7 to P21 in wild-type retinas, and significantly reduced in βII-spectrin CKO compared to controls at P30 (D). β-actin protein is shown as loading control. Co-labeling of βII-spectrin (green) with known cell-type specific markers in wild-type retinas at P30 (E). βII-spectrin protein expression (green) is significantly reduced in the OPL of βII-spectrin CKO retinas compared to controls at P30 (F). Nuclei is stained with DAPI. Scale bar shown on images.

Although several key molecules have been identified that facilitate synapse formation between photoreceptors and bipolar neurons (Cao et al., 2015; Pourhoseini et al., 2021; Sato et al., 2008), to our knowledge, none have been reported to mediate the early positioning and stabilization of bipolar dendrites to the developing synaptic layer or OPL. Moreover, the functional consequence of disrupting this early developmental event remains unclear. To answer this question, we focused on the family of spectrins, as these molecules are known to mediate key signaling events by linking the cytoskeleton to the cell membrane (Bennett & Lorenzo, 2013, 2016; Lorenzo, 2020). Although spectrins play a critical role in distinct biological functions (reviewed in (Lorenzo, 2020)), their role in synapse formation in the developing retina is unknown.

In the present study, we identified a new role for one of the members of the spectrin family, βII-spectrin in the mouse outer retina. Loss of βII-spectrin results in lamination defects whereby processes from pre- and post-synaptic neurons fail to be confined and maintained to the synaptic layer. This deficiency in synapse formation between photoreceptors and their synaptic partners leads to impaired retinal responses. Early developmental analysis revealed that βII-spectrin mediates the early dendrite positioning of rod bipolars which is required for proper circuit formation. Taken together, our findings elucidate a new developmental role for βII-spectrin in assembling retinal circuits needed for proper visual function.

## RESULTS

### βII-spectrin is localized to the developing synaptic layer

To begin to elucidate the role of βII-spectrin in the outer retina, we performed antibody staining at various developmental stages in wild-type animals. We found βII-spectrin protein localizes to the emerging OPL where protein levels increase from P7 to P30 as seen with antibody staining in **Figure 1C**. These findings were confirmed by examining βII-spectrin protein levels in whole retinas by immunoblotting (**Figure 1D**). Further examination reveals that βII-spectrin protein co-localizes to processes of distinct neuron types that project into the OPL, which includes dendrites of rod bipolars (anti-PKC), axon terminals of rod photoreceptors (anti-PSD-95), and processes of horizontal cells (anti-Calb) (**Figure 1E**). Minimal co-localization of βII-spectrin protein is seen with the axon terminals of cone photoreceptors (anti-CAR) and the dendrites of cone bipolars (anti-Scgn). Thus, these findings show that βII-spectrin is localized to the developing synaptic layer and overlaps with processes from pre- and post-synaptic neurons in the outer retina.

Next, we disrupted βII-spectrin function using a published floxed allele for βII-spectrin (*βII-spectrin*^*flox/flox*^) (Galiano et al., 2012) since complete germline knockout mice of βII-spectrin are embryonically lethal (Tang et al., 2003). To conditionally remove βII-spectrin throughout the retina, we crossed *βII-spectrin*^*flox/flox*^ animals to a *Chx10-Cre*^*+/tg*^ (Rowan & Cepko, 2004) transgenic mouse line. We refer to this cross as βII-spectrin CKO. Immunoblotting showed an overall decrease of βII-spectrin protein levels in whole retinas compared to controls (**Figure 1D**), and antibody staining confirmed knockdown of βII-spectrin within the OPL (**Figure 1F**). These findings support knockdown of βII-spectrin in the mouse outer retina using βII-spectrin CKO transgenic animals.

### Disruption of βII-spectrin results in lamination defects in the outer retina

We then examined adult retinas from βII-spectrin CKO and controls for morphological and synaptic defects. Nuclei staining with DAPI at P30 revealed no gross morphological defects in the overall retinal structure of βII-spectrin CKO compared to controls (**Figure 2A**). Measurements of the layer thickness of the ONL and INL revealed no significant difference between controls and βII-spectrin CKO (**Figure 2A**). However, the OPL was significantly thinner by 82% in βII-spectrin CKO compared to controls with several displaced nuclei found within the synaptic layer as depicted by yellow arrows in **Figure 2A**. As loss of βII-spectrin does not disrupt the overall structure of the retina, we then used several known markers of pre- and post-synaptic neurons to assess for synaptic defects. By P30, synapse formation is largely complete in the outer retina and neuronal processes from pre- and post-synaptic neurons are confined to the OPL as seen in controls in **Figure 2B**. We found that disruption of βII-spectrin leads to numerous processes (or sprouts) of horizontal cells extending beyond the OPL into the ONL (**Figure 2B**). Controls displayed an average of 0.07 ± 0.12 sprouts per 100μm length of OPL, whereas βII-spectrin CKO showed 7.70 ± 3.28 sprouts per 100μm length of OPL (**Figure 2B**). Similarly, the dendrites of rod bipolars also failed to be confined to the OPL and instead misprojected into the ONL. The average number of rod bipolar sprouts in βII-spectrin CKO was 5.83 ± 3.21 per 100μm length of OPL, whereas controls displayed no detectable sprouts (**Figure 2B**). However, very few sprouts from the dendrites of cone bipolars were observed in βII-spectrin CKO (1.90 ± 0.67 sprouts per 100μm length of OPL) as shown in **Supplemental Figure 1A-B**. These results indicate βII-spectrin is required for positioning and maintaining processes from post-synaptic neurons such as horizontal cells and rod bipolars but not cone bipolars in the OPL.

**Figure 2:**
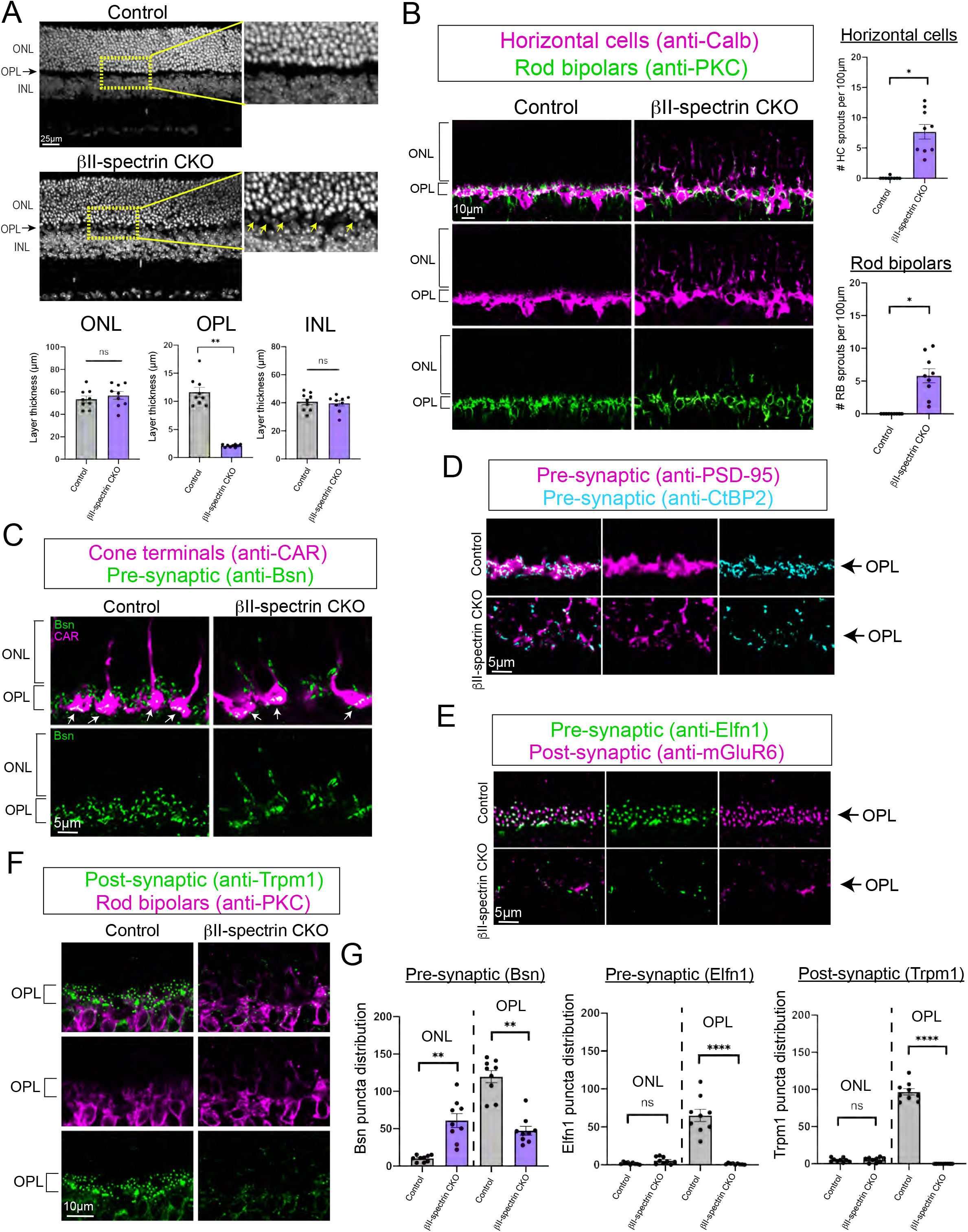
Loss of βII-spectrin results in lamination defects in the outer retina (A-G). Retinal sections of control and βII-spectrin CKO stained with DAPI reveal no gross morphological defects (A). Close examination of the OPL show several displaced nuclei in βII-spectrin CKO as depicted by the yellow arrows. Insets are zoomed images of yellow dotted boxed region. Measurements of the layer thickness show no significant difference between the ONL and INL but a significant decrease in the OPL of βII-spectrin CKO compared to controls (A). Disruption of βII-spectrin results in lamination defects were processes from horizontal cells (anti-Calb, magenta) and dendrites of rod bipolars (anti-PKC, green) failed to be confined t the OPL and instead sprout into the ONL (B). Quantification of the total number of sprouts observed in controls and βII-spectrin CKO (B). Bassoon (anti-Bsn, green) is localized to the OPL in controls but reduced in the OPL and ectopically expressed in the ONL of βII-spectrin CKO animals at P30 (C). Cone terminals (anti-CAR, magenta) are normally positioned in the OPL and express Bassoon in both controls and βII-spectrin CKO as depicted by white arrows (C). Expression and localization of pre-synaptic PSD-95 (magenta) and CtBP2 (cyan) are also affected in βII-spectrin CKO compared to controls (D). Loss of pre-synaptic Elfn1 (green) and post-synaptic mGluR6 (magenta) protein expression is seen in the OPL due to disruption of βII-spectrin (E). Post-synaptic Trpm1 (green) forms a puncta-like structure along the dendrites of rod bipolars in controls but is significantly reduced in βII-spectrin CKO (F). Quantification and localization of pre- and post-synaptic markers (i.e. Bsn, Elfn1, Trpm1) at P30 (G). The number of Bassoon puncta (∼0.6µm in size) is significantly reduced in the OPL and increased in the ONL due to loss of βII-spectrin. Pre-synaptic Elfn1 puncta (∼0.6µm in size) and post-synaptic Trpm1 puncta (∼0.6µm in size) are significantly reduced in the OPL in βII-spectrin CKO compared to controls. Images are shown as confocal sections. Data is represented as mean values ± SEM. Statistical significance determined by an unpaired two-tailed Student’s t-test. ns p>0.05, *p<0.05, **p<0.01, ****p<0.0001. Scale bar shown for each figure.

We next addressed the role of βII-spectrin on pre-synaptic neurons by examining the position of photoreceptor terminals using antibodies against Cone Arrestin (CAR) for cone terminals and PSD-95 for rod terminals. PSD-95 is expressed in both rod and cone terminals, however higher levels of protein expression are found in rods compared to cones (Sarin et al., 2018). We found the overall number of cone terminals in the OPL was not statistically different between βII-spectrin CKO (10.51 ± 1.51 cone terminals) and controls (9.87 ± 0.53 cone terminals) at P30. See **Figure 2C** and **Supplemental Figure 1B**. However, examination of rod terminals with PSD-95 staining revealed a shift in protein localization where expression becomes more diffuse (i.e. less in the OPL and ectopic puncta-like expression in the ONL) (**Figure 2D**; **Supplemental Figure 1A**). Ectopic PSD-95 protein expression in the ONL has been associated with rod retraction and loss of synaptic connectivity (Samuel et al., 2014). Thus, we set out to further examine photoreceptor synapses in βII-spectrin CKO using available antibodies.

### βII-spectrin is required for rod synaptic connectivity and retinal function

We examined protein localization of pre-synaptic markers such as Bassoon (Bsn) and CtBP2 that are both expressed in rod and cone terminals (Pourhoseini et al., 2021). We found Bsn and CtBP2 protein expression is reduced in the OPL and ectopically expressed in the ONL with disruption of βII-spectrin (**Figure 2C-D**). Bsn and CtBP2 protein form a puncta-like structure that is roughly 0.6μm and 1.0μm in diameter, respectively. The total number of Bsn and CtBP2 puncta were quantified using Imaris software in both controls and βII-spectrin CKO at P30 (**Figure 2G**; **Supplemental Figure 2**). Nearly 92% of total Bsn puncta were localized to the OPL in controls whereas 44% is found in the OPL in βII-spectrin CKO. A similar distribution was also observed with CtBP2 where 88% of puncta is in the OPL of controls and 7% in βII-spectrin CKO (**Supplemental Figure 1B**). However, the number of Bsn puncta within cone terminals was not statistically different between controls and βII-spectrin CKO (**Supplemental Figure 1A**). Next, we examined the expression of Elfn1 which is expressed in rod photoreceptors and required for selective wiring of rods to their synaptic partners (Cao et al., 2015). Interestingly, we found Elfn1 protein expression mostly absent in βII-spectrin CKO compared to controls (**Figure 2E,G**). Similarly, post-synaptic protein expression of Trpm1 and mGluR6 are also reduced due to loss of βII-spectrin (**Figure 2E,F**) with quantification of Trpm1 puncta shown in **Figure 2G**. Together, these results demonstrate βII-spectrin is required for proper localization and expression of synaptic proteins implicated in rod synaptic connectivity.

To further address if changes in synaptic protein expression translate to rod connectivity defects, we performed transmission electron microscopy (TEM) to examine the rod synaptic structure in both controls and βII-spectrin CKO at P30. Consistent with our synaptic protein findings, we found an overall reduction in the number of rod terminals localized to the OPL with several mispositioned in the ONL (**Figure 3A-B**). For controls, we analyzed a total of 56 TEM sections from three different animals and observed 1,555 rod terminals in the OPL. For βII-spectrin CKO, we analyzed 63 TEM sections from three mice and found only 296 rod terminals in the OPL (**Figure 3B**). Moreover, we classified rod terminals into four categories: (1) empty – no visible processes, (2) monads – only one invaginating process, (3) dyads – two invaginating processes, (4) triads – two invaginating processes from horizontal cells and one from a ON bipolar dendrite. Our analysis revealed an increase in the number of empty rod terminals and a decrease in the number of triads in the OPL of βII-spectrin CKO compared to controls (**Figure 3B**). These data suggest βII-spectrin is required for positioning rod terminals to the OPL.

**Figure 3:**
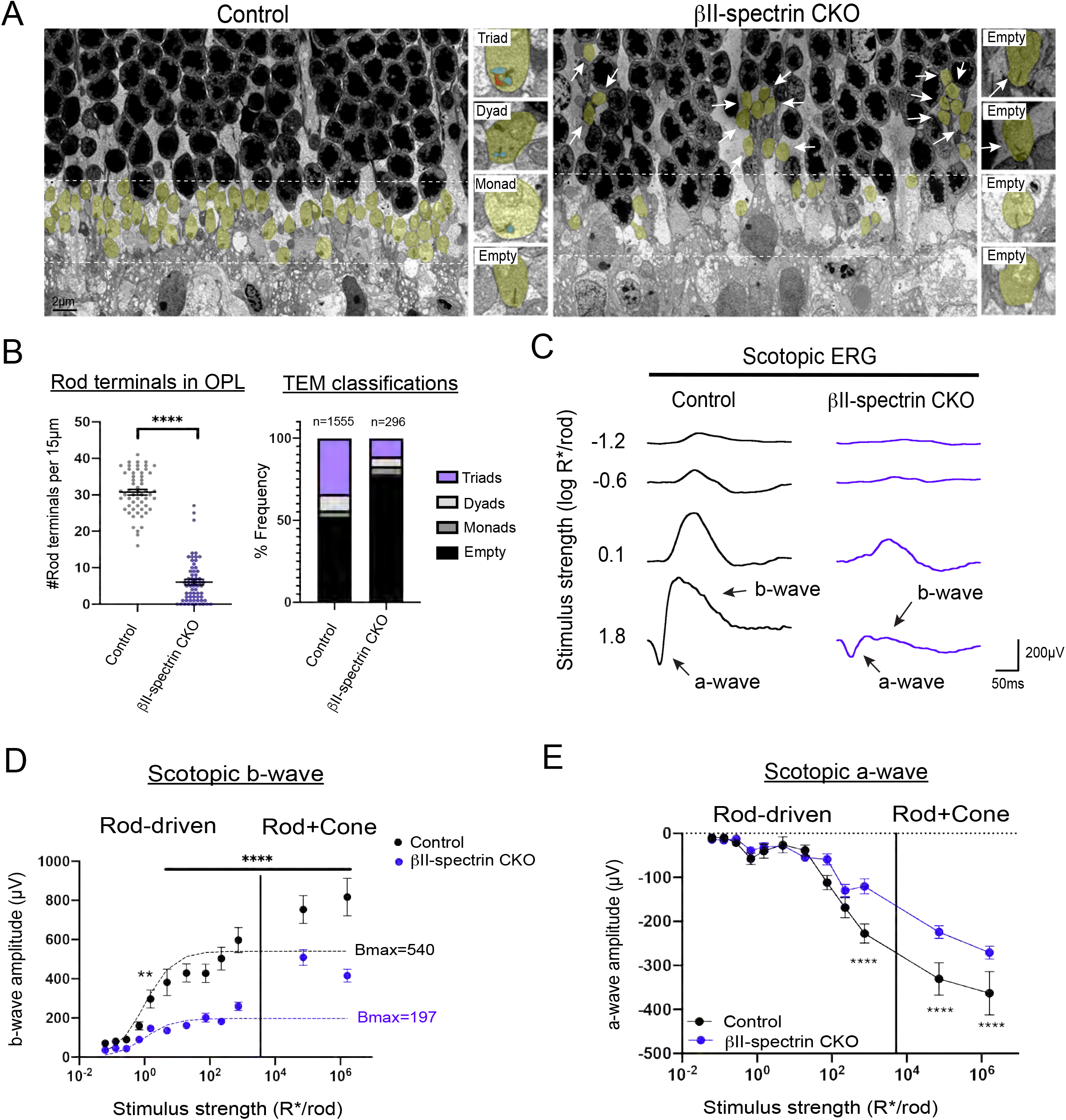
Synaptic defects and abnormal retinal responses in βII-spectrin CKO (A-E). Transmission Electron Microscopy (TEM) reveals rod terminals (yellow) are not localized to the OPL (white dotted lines) but mispositioned in the ONL (white arrows) due to loss of βII-spectrin (A). The number of rod terminals in the OPL were counted in both controls and βII-spectrin CKO (B). Data is represented as mean values ± SEM. Statistical significance determined by an unpaired two-tailed Student’s t-test. ****p<0.0001. Rod terminals were classified as either triad, dyad, monad, or empty (B). A total of 1555 rod terminals were found in the OPL and analyzed in controls; however, only 296 rod terminals were observed in the OPL of βII-spectrin CKO (B). The frequency of triads, dyads, monads, and empty in controls and βII-spectrin CKO revealed an increase in the number of empty rod terminals due to disruption of βII-spectrin (B). Individual electroretinogram (ERG) traces from controls (black line) and βII-spectrin CKO (blue line) are shown at different scotopic or rod-driven stimulus intensities (C). Scotopic b-wave amplitudes is significantly reduced in βII-spectrin CKO (blue line, n=10 from five mice) compared to controls (black line, n=8 from four mice) (D). Data points within the rod operative range are fitted with a hyperbolic saturating curve using the Naka-Rushton equation. Stimulus response plot of scotopic a-wave amplitudes of dark-adapted controls (black line, n=8 from four mice) and βII-spectrin CKO (blue line, n=10 from five mice) (E). Statistical significance determined by Holm-Sidak method for multiple comparisons. **p<0.005, ****p<0.0001.

We next asked if the rod synaptic defects observed in βII-spectrin CKO impairs retinal function by performing *in vivo* full-field electroretinograms (ERG) in dark-adapted mice. Mice were exposed to flashes of light of varying intensities to measure rod-driven responses or scotopic ERG. Cone-driven responses or photopic ERG were elicited using a paired-flash protocol as described in (Abd-El-Barr et al., 2009; Pourhoseini et al., 2021). Individual ERG traces of controls (black line) and βII-spectrin CKO (blue line) at different stimulus intensities are shown in **Figure 3C**. The a-wave represents photoreceptor responses whereas the b-wave is the downstream synaptic response from both outer and inner retinal neurons (Abd-El-Barr et al., 2009). Our results show an overall reduction in b-wave responses but not a-wave responses under scotopic conditions due to disruption of βII-spectrin. To confirm these findings, we plotted b-wave responses fitted with the Naka-Rushton equation and calculated the maximum rod-driven b-wave response (i.e. Bmax). We found controls had a Bmax value of 540μV (black dotted line) whereas βII-spectrin CKO had a Bmax of 197μV (blue dotted line) as shown in **Figure 3D**. Although scotopic b-wave responses are significantly reduced in βII-spectrin CKO compared to controls, the time-to-peak or implicit time is not affected (**Supplemental Figure 3A**). However, rod-driven or scotopic a-wave responses in dark-adapted mice were not statistically different between controls and βII-spectrin CKO at low flashes of light but were significant at higher flashes of light when both rods and cones are activated (**Figure 3E**). To further investigate cone-driven or photopic responses, we isolated the cone a-wave and b-wave and found no statistical difference between controls and βII-spectrin CKO using an unpaired Student’s t-test (p<0.05) as shown in **Supplemental Figure 3B**. Together, these data show impaired rod-driven downstream responses or scotopic b-wave in βII-spectrin CKO compared to controls, which support the synaptic connectivity defects observed with loss of βII-spectrin.

### Temporal requirements of βII-spectrin in the outer retina

We then asked when is βII-spectrin required for synaptic connectivity in the outer retina. To answer this question, we performed a developmental analysis at early stages of synaptogenesis (i.e. P9, P11, P13) in βII-spectrin CKO animals. We focused our analysis on rod bipolars and horizontal cells as these retinal neurons show significant lamination defects at adult stages (**Figure 2B**). We found that by P9 the dendrites of rod bipolars extend into the OPL and form one continuous layer as seen in controls (**Figure 4A**). However, we observed gaps or areas where the dendrites of rod bipolars failed to extend in the OPL (yellow arrow) and instead sprout into the ONL (white arrows) due to loss of βII-spectrin (**Figure 4A**). The total number of rod bipolar sprouts continued to increase from P9 to P13 as shown in **Figure 4B**. Interestingly, we did not observe gaps or sprouts of processes from horizontal cells at P9 (**Figure 4A**). However, the number of horizontal cell sprouts did increase from P11 to P13 in βII-spectrin CKO compared to controls (**Figure 4A-B**). These findings suggest that the improper positioning of the dendrites of rod bipolars at early stages (i.e. P9) may lead to the subsequent lamination defects seen in horizontal cells. We also examined the position of cone terminals (anti-S-opsin), dendrites of cone bipolars (anti-Scgn), and rods (anti-Rhodopsin) at P9 (**Figure 4C**). Although we do see gaps or defects in dendrite extension of rod bipolars (anti-PKC) and loss of pre-synaptic protein expression (anti-PSD-95) in βII-spectin CKO, we do not observe defects in the positioning of other retinal cell types (**Figure 4C**). These findings further support the model that βII-spectrin is required early on to position the dendrites of rod bipolars to the emerging OPL.

**Figure 4:**
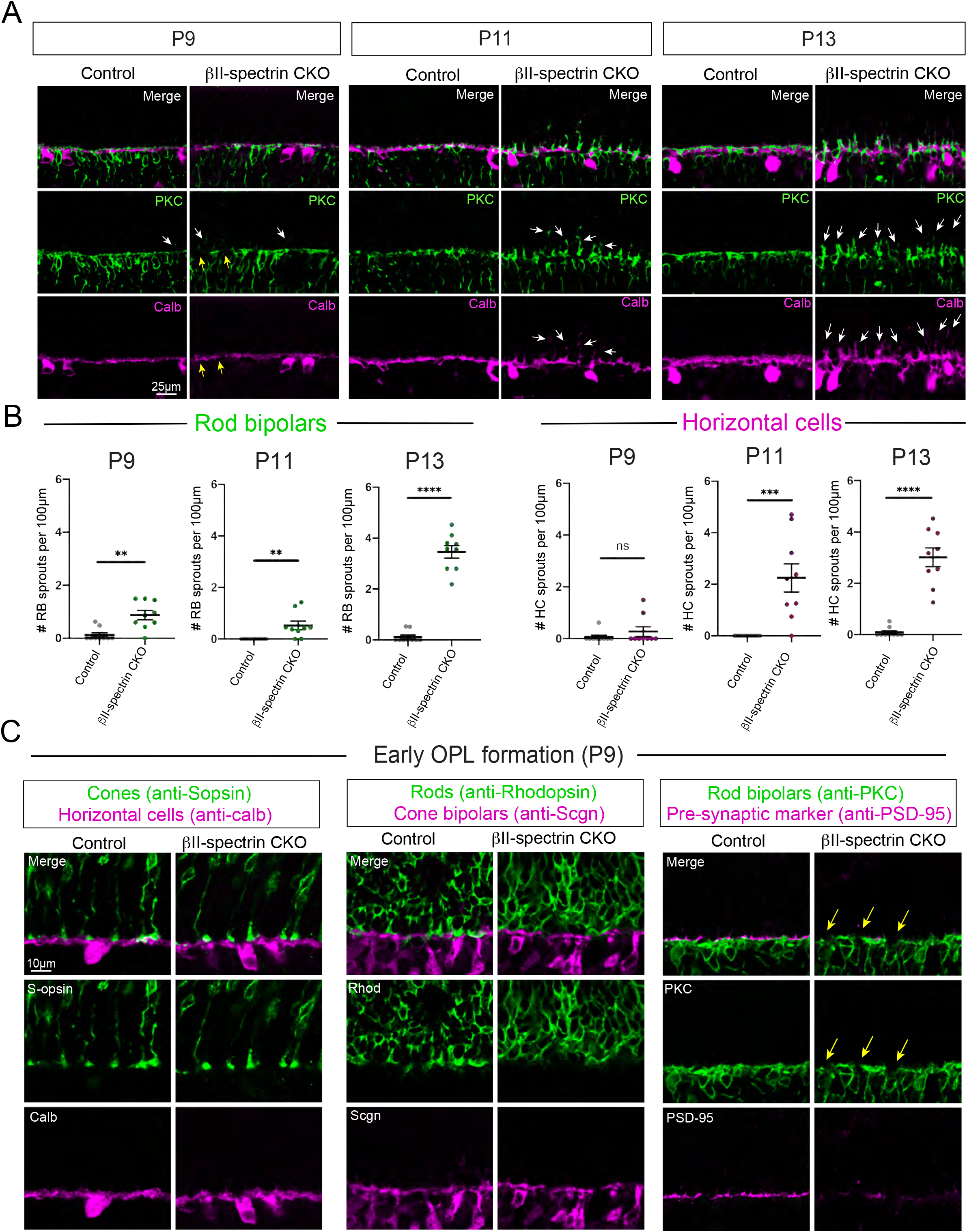
Developmental analysis of βII-spectrin CKO (A-C). At P9, dendrites of rod bipolars (green, anti-PKC) and processes from horizontal cells (magenta, anti-calb) form one continuous laminated structure in controls (A). However, several mispositioned dendrites of rod bipolars such as gaps (yellow arrows) and sprouts (white arrows) are observed in βII-spectrin CKO. Mispositioned processes or sprouts from horizontal cells (white arrows) are not detected until P11 in βII-spectrin CKO and continue to P13 (A). Quantification of the total number of rod bipolar and horizontal cell sprouts per 100µm length of OPL across the different time points (B). Few sprouts are observed in controls compared to βII-spectrin CKO from P9 to P13 (A,B). Statistical significance determined by an unpaired two-tailed Student’s t-test. **p< 0.005, ****p<0.0001. Processes from other pre- and post-synaptic neurons are normally positioned to the nascent OPL at P9 (C). Cone terminals (green, anti-Sopsin) and processes from horizontal cells (magenta, anti-calb) are localized to the developing OPL in both controls and bII-spectrin CKO. Rods (green, anti-Rhodopsin) and dendrites of cone bipolars (magenta, anti-Scgn) are largely confined to the OPL at P9 in controls and bII-spectrin CKO. Dendrites of rod bipolars (green, anti-PKC) begin to display gaps and loss of pre-synaptic protein expression (magenta, anti-PSD-95) as depicted by yellow arrows due to loss of βII-spectrin. Scale bar shown on figures.

## DISCUSSION

In the present study, we uncovered a new role of βII-spectrin in assembling neural circuits in the mouse outer retina. We found βII-spectrin positions and maintains the processes of pre- and post-synaptic neurons to the developing synaptic layer or OPL. Loss of βII-spectrin results in lamination defects where processes fail to be confined to the OPL resulting in impaired synaptic connectivity and abnormal retinal responses. These findings highlight a new key player in neural circuit formation required for normal visual function.

### Spectrins in the developing nervous system

Spectrins play an essential role in the central nervous system. They are known to bind to cell adhesion molecules, receptors, ion channels, and other scaffolding proteins (Bennett & Lorenzo, 2013, 2016; Lorenzo, 2020). Individuals with mutations in SPTBN2, the gene encoding for βIII-spectrin has been associated with spinocerebellar ataxia type 5 in humans (Ikeda et al., 2006). Mice with loss of βIII-spectrin display defects in dendrite morphology of Purkinje cells and cerebellar degeneration (Perkins et al., 2010). Human mutations in SPTBN1, gene encoding for βII-spectrin display mild to severe intellectual disabilities, language and motor deficits, seizures, and behavioral abnormalities (Cousin et al., 2021). Similarly, mice that lack βII-spectrin in neural progenitors display motor coordination deficits, multiple seizures, reduced long-range axonal connectivity, and early postnatal lethality (Lorenzo et al., 2019). The widespread neurological defects observed with loss of spectrins suggest that they have distinct functions in setting up different neural circuits.

Our findings uncovered a new key molecule that is required for the early positioning and stabilization of late-born neurons to the emerging synaptic layer. Although dendrite extension of bipolar neurons into the OPL has been well-documented (Morgan et al., 2006), the significance of this process on synapse formation had been relatively unknown. We found loss of βII-spectrin disrupts the normal positioning of the dendrites of rod bipolar at early stages and this leads to defects in synaptic connectivity and impaired retinal responses in the adult. Throughout the central nervous system, neurons are integrated into a nascent neural circuit at various stages during development (Faux et al., 2012). Defects in neuron migration, targeting, and connectivity are often linked to neurodevelopmental disorders such as intellectual disability, epilepsy, and autism spectrum disorder (Bozzi et al., 2012; Moon & Wynshaw-Boris, 2013; Pan et al., 2019). However, disseminating the mechanisms involved in neural circuit assembly has remained challenging. Several key molecules including spectrins are known to play multiple roles throughout development (Lorenzo, 2020). Spectrins are known to mediate early roles in neurodevelopment such as cell migration, neuron morphology, and axon outgrowth (Cousin et al., 2021; Lorenzo, 2020; Lorenzo et al., 2019). Thus, the retina is an excellent model where we can take advantage of the different experimental approaches that can tease apart the early and later temporal requirements of spectrins to elucidate their multiple functions.

### βII-spectrin function in the outer retina

The mouse retina is an excellent system to uncover new developmental mechanisms of neural circuit assembly. The various transgenic mouse lines and tools that are readily available in the retina allows us to disrupt gene function in both a cell-type and temporal manner. Germline knockouts of βII-spectrin are embryonically lethal (Tang et al., 2003). Thus, we were able to bypass the early requirements of βII-spectrin using retinal-specific cre lines that turns on at later stages in development (Rowan & Cepko, 2004). This allowed us to uncover a new role of βII-spectrin in assembling neural circuits that occurs at later developmental time points.

Our developmental analysis showed that βII-spectrin is required early on to position the dendrites of rod bipolars to the emerging synaptic layer. As lamination defects are often seen in mouse mutants with synaptic connectivity defects (Dick et al., 2003; Ribic et al., 2014; Soto et al., 2013), this raises the question whether βII-spectrin is directly involved in remodeling the cytoskeleton or indirectly mediates synapse formation through recruitment of cell adhesion molecules. For instance, recent studies show that βII-spectrin acts on the axon terminal of cultured neurons, where loss of βII-spectrin disrupts axonal transport and inhibits axon growth (Lorenzo et al., 2019). βII-spectrin has been shown to interact with membrane lipids and this is required for dynein/dynactin axonal transport (Muresan et al., 2001). Mutations in βII-spectrin that disrupt binding with membrane lipids leads to defects in axonal transport and growth (Hyvonen et al., 1995). These data raise the possibility that βII-spectrin in rod bipolars may mediate the direct outgrowth of the dendrites into the developing synaptic layer by interacting with membrane lipids to mediate dendritic transport and growth. Another possibility is that βII-spectrin may act indirectly on synapse formation by recruiting and maintaining cell adhesion molecules to the cell surface. Studies in myelinated axons demonstrate spectrins bind to cell adhesion molecules to mediate the precise positioning of the nodes of Ranvier (Galiano et al., 2012; Rasband & Peles, 2021). One of the known binding partners of spectrins is the L1-family cell adhesion molecule, Neurofascin (Eshed et al., 2005; Ho et al., 2014; Rasband & Peles, 2021). Previous work from our lab showed Neurofascin is expressed in the developing synaptic layer and required for proper connectivity between rod photoreceptors and their synaptic targets (Pourhoseini et al., 2021). However, loss of Neurofascin does result in lamination defects as those seen with loss of βII-spectrin suggesting that there may be other binding partners in the outer retina. Future studies are needed to identify the downstream mechanism of how βII-spectrin mediates synapse formation in the developing retina.

### Assembling complex neural circuits during development

Recent work in the retina begin to shed light into the complexity of assembling neural circuits. Over the last decade, several new molecules have been identified to mediate different aspects of synapse formation and maintenance in the outer retina (Burger et al., 2021; Hoon et al., 2014; Zhang et al., 2017). However, how these molecules work together during development to mediate the precise connectivity of synaptic partners remains unclear. Studies using different mutant mouse lines begin to illustrate differences in synaptic phenotypes when different molecules are disrupted during synaptogenesis. These phenotypes can be largely grouped into three main classes: (i) defects in both retinal lamination and synaptic connectivity, (ii) defects in retinal lamination only, and (iii) defects in synaptic connectivity only (Burger et al., 2021; Zhang et al., 2017). Although our findings revealed βII-spectrin leads to defects in both retinal lamination and synaptic connectivity, the changes in protein expression reflect a later stage in synapse formation. Both Bassoon and Elfn1 are localized to the axon terminal of rod photoreceptors and loss of protein expression in mutant mice disrupts rod synaptic connectivity (Cao et al., 2015; Dick et al., 2003). Our data revealed loss of βII-spectrin leads to ectopic expression of Bassoon and CtBP2 but absence of Elfn1 and PSD-95. Previous work has shown Bassoon expression appears at early stages in synapse development (Sarin et al., 2018), whereas Elfn1 turns on at later stages and expression is reinforced after synapse formation (Cao et al., 2015). These data show how different synaptic proteins may be required at different stages during synaptogenesis. Similarly, recent work on the voltage-gated calcium channel (Ca_v_1.4) revealed that it has different developmental functions in rod photoreceptor synapses (Maddox et al., 2020). Complete knockout of the Cav1.4 channel results in retinal lamination defects and absence of rod photoreceptor synapses (Liu et al., 2013; Regus-Leidig et al., 2014; Zabouri & Haverkamp, 2013). However, a single point mutation in Ca_v_1.4 that abolishes Ca2+ influx leads to a milder phenotype where photoreceptor synapses are still largely formed but they are not functional (Maddox et al., 2020). These data suggest that the calcium channel Ca_v_1.4 may be required at early stages to structurally recruit key proteins to the synapse and at later stages to mediate functional activity. βII-spectrin may have a similar role where it functions at later developmental stages to recruit specific molecules to aid in synapse formation of the rod pathway.

Taken together, our work in the retina uncovered a new developmental role of βII-spectrin in building neural circuits. The discovery of new molecular pathways could provide insights into mechanisms of assembling neural circuits and how these could lead to aberrant wiring in individuals with neurodevelopmental disorders.

## Experimental Procedures

### Mouse strains

All mouse procedures were approved by Baylor College of Medicine Institutional Animal Care and Use Committee. *βII-spectrin*^*flox/flox*^ have been described previously (Galiano et al., 2012) and were kindly provided by Matthew Rasband, Baylor College of Medicine. To remove βII-spectrin throughout the retina, *βII-spectrin*^*flox/flox*^ mice were crossed with *Chx10-Cre*^*+/tg*^ (Rowan & Cepko, 2004) to generate βII-spectrin CKO. *βII-spectrin*^*flox/flox*^ littermates without cre served as controls for all experiments. Wild-type retinas are from CD1 mice purchased from Charles River. Both males and females were used in all experiments.

### Immunohistochemistry

Eyes were collected at various developmental time points with P0 designated as the day of birth. Whole eyes were fixed at 60 mins in 4% paraformaldehyde in PBS except for mGluR6, Trpm1, CtBP2, and Elfn1 staining which were lightly fixed at room temperature for 10 mins. Eye cups were dissected and sectioned at 20µm as previously described (Pourhoseini et al., 2021). Slides were dried overnight and washed with PBS for 10 mins twice to start antibody staining. Sectioned slides were incubated with blocking buffer (10% normal goat serum, 1% BSA, 0.5% Triton X-100 in PBS) followed by primary antibodies at 4°C overnight (**Supplemental Figure 4**). Slides were washed 3 times with PBS for 10 mins each and then incubated with secondary antibodies at 1:1000 dilution at 4°C overnight. Slides were then washed 3 times with PBS, stained with DAPI (1:1000), and then sealed with Vectashield (Vector Laboratories). For Elfn1 staining, retinal sections were pretreated with antigen retrieval reagents (R&D systems, CTS013). Images were acquired on a Zeiss LSM 800 confocal microscope and analyzed using Imaris confocal software version 9.6 (Bitplane, South Windsor, CT, USA).

### Histological quantification

For quantification, images were collected from three different retinal sections from three animals per group (n=9) as described previously (Pourhoseini et al., 2021). βII-spectrin antibody staining was performed in all retinal sections to confirm loss of βII-spectrin in βII-spectrin CKO. All confocal images were taken from the central-periphery region of the retina. Confocal images were acquired using a Zeiss LSM 800 microscope with a 40X objective and then analyzed with Imaris confocal software. Quantification of pre- and post-synaptic marker expression was performed using the Imaris Spot feature. Puncta with a diameter of 0.6µm was used to detect protein expression of Bassoon, Elfn1, and Trpm1. Puncta with a diameter of 1.0µm was used for CtBP2 protein expression. The number of puncta in the ONL was normalized to 0.01mm^2^ by computing the surface area of each individual retinal section. This was done by measuring the height and width of the ONL using DAPI staining. The number of puncta in the OPL per 100μm length was also normalized by measuring the total length of the OPL in each retinal section. Retinal layer thickness was measured in confocal sections stained with DAPI using Imaris. Statistical significance between controls and experimental groups was determined using an unpaired two-tailed Student’s t-test. All statistical analysis were performed using GraphPad Prism version 9 with p-values given in the text and figure legends.

### Transmission Electron Microscopy (TEM)

Eye cups were fixed in 3% glutaraldehyde in 4°C overnight. Tissue samples were washed with 1M sodium phosphate buffer (pH 7.3), post-fixed in 1% osmium tetroxide for 1 hour and dehydrated through a series of graded alcohol steps. Tissue samples were infiltrated (harden) with acetone and polybed 812 plastic resin and embedded in plastic block molds with 100% polybed 812. Ultra-thin sections (80nm) were cut using a Leica EMUC ultra microtome. These sections were mounted on 100 mesh copper grids and stained with 2% uranyl acetate and Reynold’s lead stain. Grids were visualized on a JEOL JEM 1230 electron microscope and images were captured on an AMTV600 digital camera. TEM analysis was performed on three different controls and three βII-spectrin CKO animals. Statistical significance was determined using an unpaired Student’s t-test.

### Electroretinography (ERG)

Scotopic ERGs were recorded bilaterally from four control animals (n=8) and five βII-spectrin CKO mice (n=10) at 6-8 weeks old. Mice were dark adapted overnight and anesthetized with a weight-based i.p. injection solution of ketamine (46 mg/ml), xylazine (9.2 mg/ml), and acepromazine (0.77 mg/ml). We performed the ERG as described in (Pourhoseini et al., 2021). Platinum electrodes were placed at the base of the tail and another between the ears to serve as ground and reference electrodes, respectively. Mice were moved into a Ganzfeld dome and remained in complete darkness for 5 minutes before initiating the experiment. Half millisecond square flashes for scotopic measurements were produced by cyan light emitting diodes of 503 nm peak wavelength. The output of the LED flashes was calibrated using a radiometer (ILT1700 International Light, MA) with a photodiode sensor and scotopic filter that provided readout in the unit of scot cd*s/m^2^. These were converted to the unit of photoisomerizations/rod (R*/rod) where 1 scot cd*s/m^2^= 581 photoisomerizations/rod/s as previously reported (Abd-El-Barr et al., 2009; Tse et al., 2015). Photopic ERG responses were elicited by using a paired-flash protocol where an initial conditioning flash (4.6 × 10^2^ R* per M cone) saturates both rods and cones 2 seconds before a probe flash (1.8 × 10^2^ R* per M cone) as described (Abd-El-Barr et al., 2009; Pourhoseini et al., 2021; Tse et al., 2015). Statistical significance was determined using the Holm-Sidak method for multiple comparisons or an unpaired Student’s t-test using the GraphPad software.

## Acknowledgements

We thank Matthew Rasband and Melanie Samuel for generously providing transgenic mouse lines, antibodies, and reagents. We also thank Ross Poché and Ching-Kang (Jason) Chen for insightful discussions throughout the development of the study. We would also like to thank Guofu Shen for help with ERG, and Ralph Nichols for technical assistance with the TEM experiments. This work was supported by the National Eye Institute (R00EY028200, R01EY033037 to EZS), an ARVO Genentech Career Development Award to EZS, Research to Prevent Blindness (RPB) Career Development Award to EZS, and a P30EY002520 and an unrestricted grant from RPB to the Department of Ophthalmology.

## Author Contributions

DGS and EZS planned and designed the experiments. DGS, CB, and JTA collected tissue for histological analyses and performed antibody staining. DGS and EZS acquired the confocal images. CB performed the histological quantification. AM conducted the TEM analysis. DGS performed the ERG recordings. DGS and EZS wrote the manuscript.

## Figure Legends

**Supplemental Figure 1: Minimal defects in the cone pathway due to loss of βII-spectrin (A-B)**. Pre-synaptic PSD-95 protein (green) is enriched in rod terminals as seen in controls (A). Redistribution of PSD-95 protein expression is observed in bII-spectrin CKO where there is loss in the OPL and ectopic expression in the ONL (A). Dendrites of cone bipolars (magenta, anti-Scgn) show minimal number of sprouts in bII-spectrin CKO compared to controls (A,B). Quantification of the total number of cone bipolar sprouts, number of cone terminals in OPL, and number of pre-synaptic Bassoon (Bsn) puncta in cone terminals (B). The number of cone terminals in the OPL and the number of Bsn puncta were not statistically different (p>0.05) between controls and βII-spectrin CKO. Statistical significance determined by an unpaired two-tailed Student’s t-test. **p< 0.005. Scale bar, 10µm.

**Supplemental Figure 2: Ectopic expression of pre-synaptic CtBP2 in βII-spectrin CKO (A)**. The total number of CtBP2 puncta (1.0μm in size) in the ONL and OPL were detected using the Imaris software. Controls show minimal CtBP2 puncta in the ONL and higher number in the OPL, whereas βII-spectrin CKO show a higher number in the ONL compared to the OPL. Statistical significance determined by an unpaired two-tailed Student’s t-test. **p< 0.005.

**Supplemental Figure 3: Scotopic and photopic ERG responses of βII-spectrin CKO animals (A-B)**. Rod-driven implicit times are shown for controls (black line, n=8 from four mice) and βII-spectrin CKO (blue line, n=10 from five mice) (A). Isolated amplitudes of the cone b-wave and a-wave using the paired flash method (B). Data represented as mean values ± SD. Statistical significance determined by an unpaired two-tailed Student’s t-test (p>0.05).

