## Supplemental Figures for "βII-spectrin is required for synaptic positioning during retinal development"

Supplemental Figure 1 related to Figure 2: Cone pathway

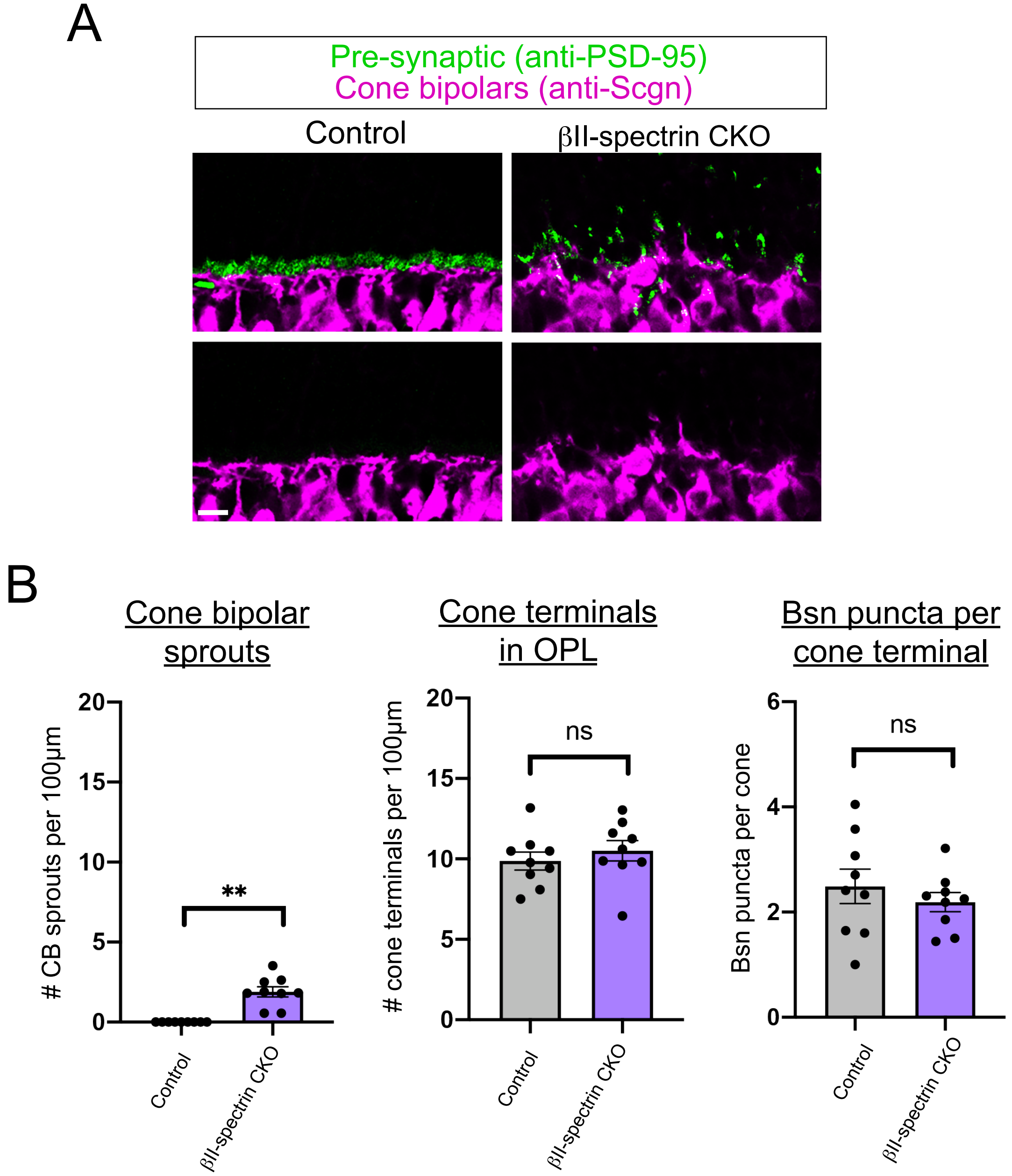

Supplemental Figure 2: CtBP2 quantification

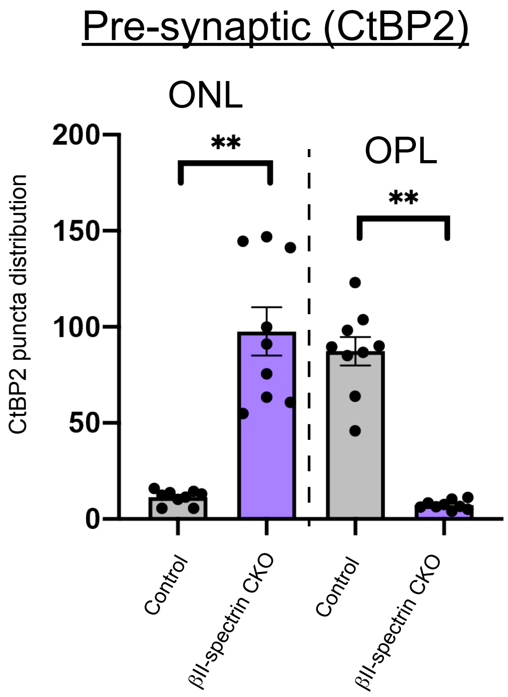

Supplemental Figure 3 related to Figure 3: ERG responses

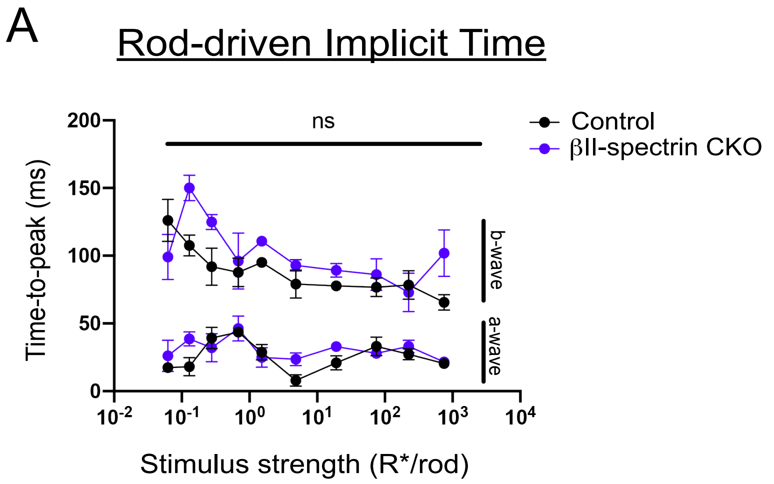


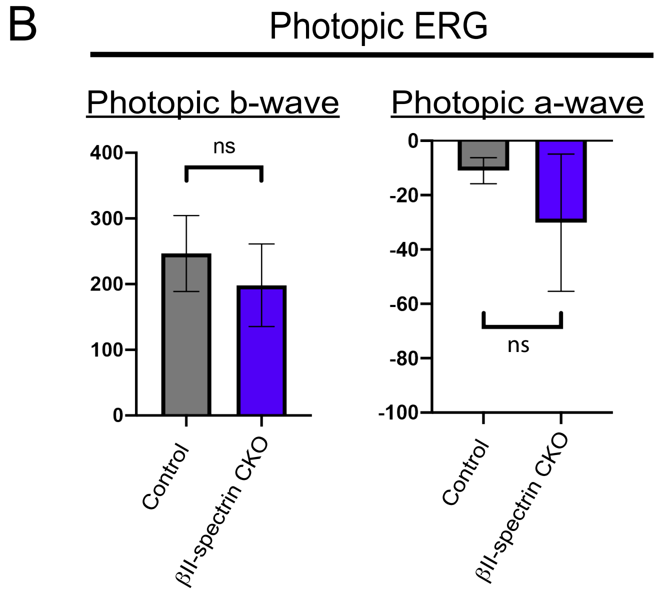

Supplemental Figure 4: Table of antibodies

| Antibody name | Labeling specificity | Source | Dilution |
|---|---|---|---|
| Mouse anti-βII-spectrin | βII-spectrin | BD Biosciences Cat# 612562, RRID:AB_399853 | 1:250 |
| Rabbit anti-Cone Arrestin (CAR) | Cone Photoreceptors | Millipore Cat# AB15282, RRID:AB_1163387 | 1:500 |
| Mouse monoclonal anti-Bassoon | Presynaptic photoreceptor terminals | Enzo Life Sciences Cat# VAM-PS003F, RRID:AB_1659573 | 1:500 |
| Rabbit anti-Calbindin (Calb) | Horizontal cells, amacrine cells, retinal ganglion cells | Swant Cat# CB38, RRID:AB_10000340 | 1:2000 |
| Mouse monoclonal anti-Protein Kinase C (PKC) | Rod bipolars | Abcam Cat# ab31, RRID:AB_303507 | 1:500 |
| Mouse polyclonal anti-PSD-95 | Photoreceptor terminals (highly expressed in rods compared to cones) | Thermo Fisher Scientific Cat# MA1-046, RRID:AB_2092361 | 1:500 |
| Rabbit polyclonal anti-metabotropic Glutamate Receptor 6 (mGluR6 ) | ON bipolar neurons | Gift from Larry Zipursky; (Sarin et al., 2018) | 1:500 |
| Rabbit polyclonal anti-Secretagogin (Scgn) | Cone bipolar cells | BioVendor Laboratory Medicine Cat# RD181120100, RRID:AB_2034060 | 1:1000 |
| Mouse monoclonal anti-CtBP2 | Photoreceptor Terminals | BD Biosciences Cat# 612044, RRID:AB_399431 | 1:500 |
| Sheep anti-Trpm1 | Post synaptic ON bipolar | Gift from Kirill Martemyanov; (Cao et al., 2015) | 1:500 |
| Rabbit anti-Elfn1 305-320 | Rod photorecptor terminals | Gift from Kirill Martemyanov; (Cao et al., 2015) | 1:100 |
